# Reproducible acquisition, management, and meta-analysis of nucleotide sequence (meta)data using q2-fondue

**DOI:** 10.1101/2022.03.22.485322

**Authors:** Michal Ziemski, Anja Adamov, Lina Kim, Lena Flörl, Nicholas A. Bokulich

## Abstract

The volume of public nucleotide sequence data has blossomed over the past two decades, enabling novel discoveries via re-analysis, meta-analyses, and comparative studies for uncovering general biological trends. However, reproducible re-use and management of sequence datasets remains a challenge. We created the software plugin *q2-fondue* to enable user-friendly acquisition, re-use, and management of public nucleotide sequence (meta)data while adhering to open data principles. The software allows fully provenance-tracked programmatic access to and management of data from the Sequence Read Archive (SRA). Sequence data and accompanying metadata retrieved with *q2-fondue* follow a validated format, which is interoperable with the QIIME 2 ecosystem and its multiple user interfaces. To highlight the manifold capabilities of *q2-fondue*, we present several demonstration analyses using amplicon, whole genome, and shotgun metagenome datasets. These use cases demonstrate how *q2-fondue* increases analysis reproducibility and transparency from data download to final visualizations by including source details in the integrated provenance graph. We believe *q2-fondue* will lower existing barriers to comparative analyses of nucleotide sequence data, enabling more transparent, open, and reproducible conduct of meta-analyses. *q2-fondue* is a Python 3 package released under the BSD 3-clause license at https://github.com/bokulich-lab/q2-fondue.

## Introduction

The increasing volume of publicly available nucleotide sequence data is driving a revolution in the life sciences, by enabling comparative studies to discover generalizable trends that are often inaccessible or underpowered in an individual study. Individual studies addressing similar biological questions can differ in many technical aspects, including (but not limited to) specific experimental design, employed sequencing technologies, definitions of the examined target variables, and selection of potential covariates influencing the target. These inter-study variations can make individual study results biased (Serghiou et al., 2016) and even contradictory to one another (Ioannidis & Trikalinos, 2005). Meta-analysis allows the synthesis of findings from individual studies to reach a more complete understanding: identifying consistent and reproducible signatures across studies, and resolving causes of variation among study results (Gurevitch et al., 2018).

Meta-analyses of nucleotide sequencing-based studies have intensified within the past decade (see Figure 1), given the high potential of these data for reuse in comparative analyses. Meta-analyses of genome-wide association data have expanded our knowledge of human polygenic disorders and quantitative traits (Panagiotou et al., 2013). Comparative genomics has given insights into vertebrate genome evolution (Meadows & Lindblad-Toh, 2017) and the processes of genome function, speciation, selection and adaptation (Alföldi & Lindblad-Toh, 2013). Comparative analyses of global microbiome datasets have driven deepening insight into spatiotemporal and biogeographic variation in Earth’s microbial diversity (Tara Oceans Consortium Coordinators et al., 2015; Thompson et al., 2017). Re-analysis of published genome and metagenome data have enabled discovery of novel and candidate microbial clades, as in the Genome Taxonomy Database (Parks et al., 2017), and highlighted the abundance (Lloyd et al., 2018) and ecosystem-impact (Zamkovaya et al., 2021) of uncultured microbes, also known as “microbial dark matter”.

**Figure 1.**
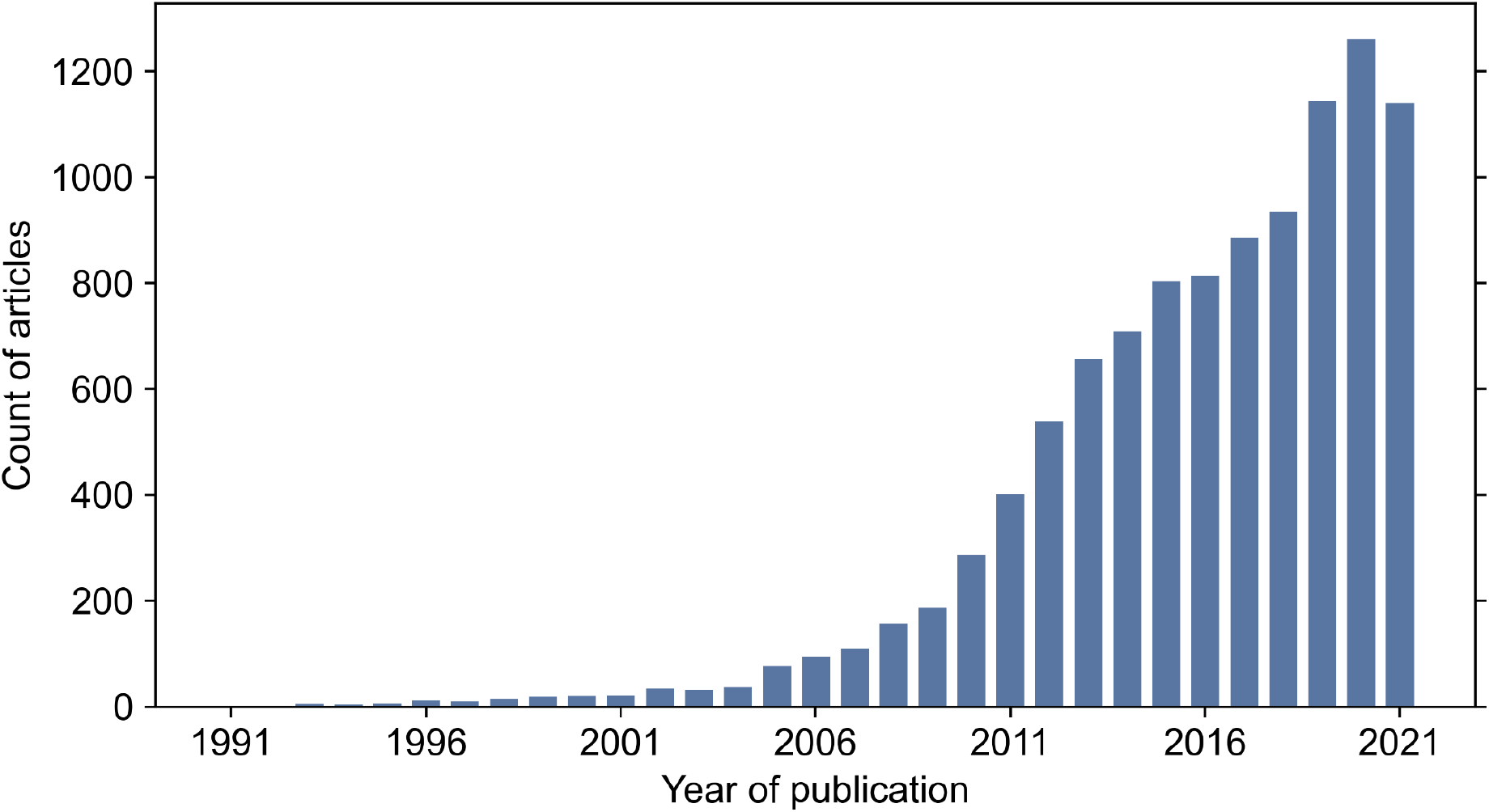
Increasing trend of sequencing-based meta-analyses over the past 30 years. Displayed article counts were retrieved from PubMed on 2022/02/21 with the search query “(meta-analysis) AND ((omics) OR (genom*) OR (microbio*) OR (transcriptom*))” and a requirement for the article type to be a meta-analysis.

Since meta-analyses can only be conducted if the original study data are publicly available, the recent increase of meta-analyses can be partly attributed to the ongoing open-science efforts of making sequencing data and accompanying metadata standardized and publicly available (Berman et al., 2014; McNutt et al., 2016; “The Path to Open Data,” 2019; Yilmaz et al., 2011). The Sequence Read Archive (SRA), established as part of the International Nucleotide Sequence Database Collaboration (INSDC) by the National Center for Biotechnology Information (NCBI), enables such free access to sequence data (Kodama et al., 2012; Leinonen, Sugawara, et al., 2011), including sequences stored on ENA (Leinonen, Akhtar, et al., 2011) and DDBJ (Mashima et al., 2017). Since its creation in 2009, the SRA has gathered data at the petabyte scale and continues to scale its infrastructure to ensure efficient data storage and retrieval (Katz et al., 2021).

A selection of tools to programmatically access data from the SRA has recently emerged. NCBI offers the *sra-tools* command-line toolkit (Leinonen, Sugawara, et al., 2011) for downloading and interacting with raw sequence data. The *entrezpy* Python library (Buchmann & Holmes, 2019) aids automating the data download from NCBI’s Entrez databases by providing abstract classes allowing custom implementations. *pysradb* (Choudhary, 2019) makes use of the curated metadata database available through the SRAdb project (Zhu et al., 2013) to download data from the SRA, and *grabseqs* enables fetching of data from not only the SRA but also MG-RAST (Meyer et al., 2008) and iMicrobe (Youens-Clark et al., 2019). However, wider application of these tools in comparative analyses is hindered by several challenges, including technical complication and a steep learning curve, and the difficulty of tracking data provenance necessary for reproducibility and traceability in large comparative studies.

In order to provide consistent and reliable findings, meta-analyses must follow Findable, Accessible, Interoperable and Reusable (FAIR) Guiding Principles (Wilkinson et al., 2016). To this end, meta-analyses should be performed in a reproducible manner, making use of consistent workflows while keeping track of all the performed data retrieval and analysis steps. Despite increasingly facilitated access to sequencing data, reproducibility and provenance of primary and secondary studies remain challenging (Amann et al., 2019; Baker, 2016; Huang & Gottardo, 2013; Kim et al., 2018; Reichman et al., 2011). Here we introduce an open-source software package *q2-fondue* (**F**unctions for reproducibly **O**btaining and **N**ormalizing **D**ata re**u**sed from **E**lsewhere) to expedite the initial acquisition of data from the SRA, while offering complete provenance tracking. *q2-fondue* simplifies retrieval of sequencing data and accompanying metadata in a validated and standardized format interoperable with the QIIME 2 ecosystem (Bolyen et al., 2019). By allowing access through multiple QIIME 2 user interfaces, it can be employed by users of different computational capabilities.

Here we describe the *q2-fondue* software package and subsequently demonstrate its use in comparative analyses of marker-gene, genome, and metagenome sequencing studies. We anticipate *q2-fondue* will lower existing barriers to comparative analyses of nucleotide sequence data, facilitating more transparent, open, and reproducible conduct of meta-analyses.

## Methods

### Implementation and overview

*q2-fondue* is an open source Python 3 package released under the BSD 3-clause license. It can be installed in a conda environment on any UNIX-based system as described in the installation instructions provided on the package website (https://github.com/bokulich-lab/q2-fondue). *q2-fondue* has been implemented as a QIIME 2 plugin, allowing use of QIIME 2’s integrated data provenance tracking system, multiple user interfaces, and streamlined interoperability with downstream sequence analysis plugins.

An overview of q2-fondue is shown in Figure 2. Two separate q2-fondue actions allow easy access to the SRA database: *get-sequences* and *get-metadata* fetch per-sample sequence data and corresponding metadata (e.g., sample and run information), respectively. The *get-all* pipeline wraps both of these actions to simultaneously retrieve sequences and metadata for a list of SRA accessions. These three actions, *get-sequences, get-metadata* and *get-all*, require a single input file containing accession IDs of SRA runs or BioProjects to fetch. SRA run IDs allow direct interaction with the SRA databases, while BioProject IDs are first translated into corresponding run IDs using a chain of E-Direct utilities (Kans, 2013). An *E-Search* query is executed to look up provided IDs in the BioProject database, followed by an *E-Link* query finalized by an *E-Fetch* query to retrieve the linked run IDs.

**Figure 2.**
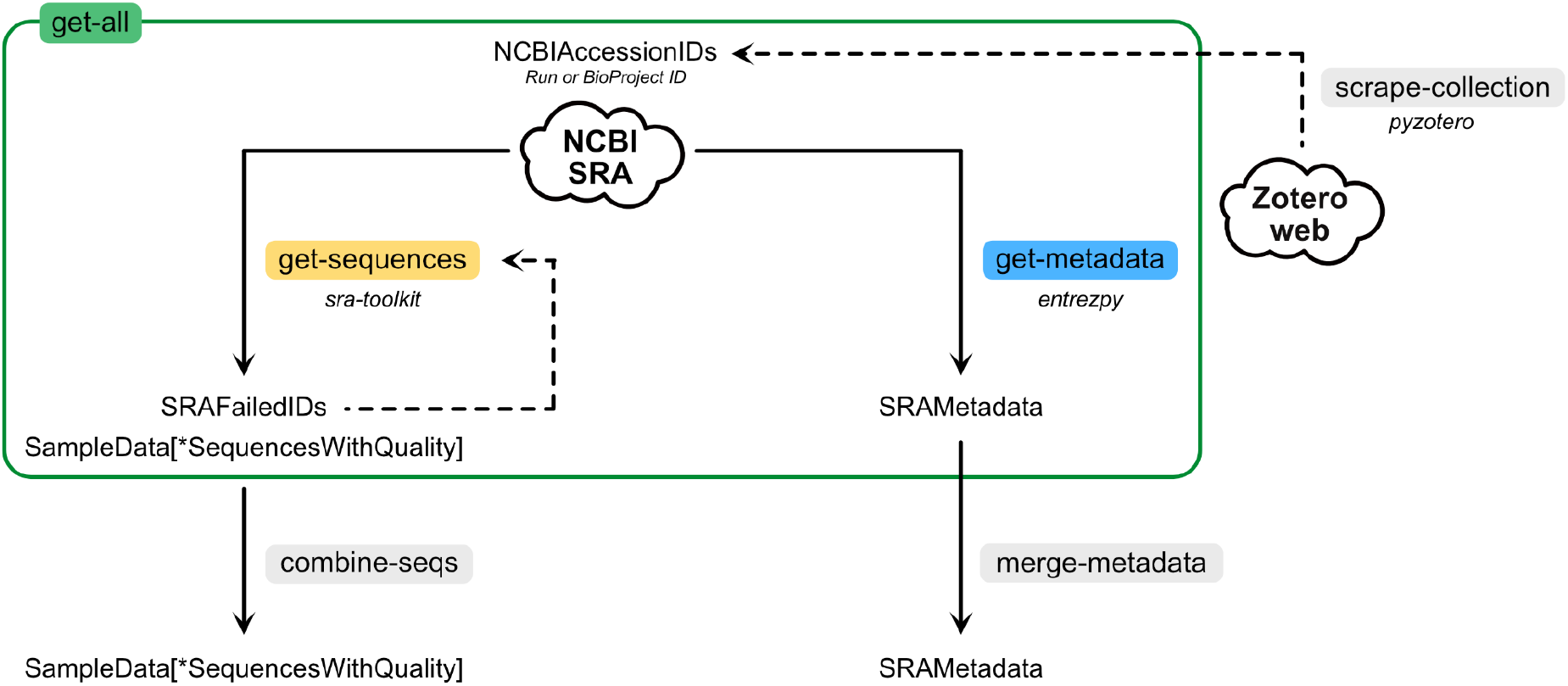
Overview of *q2-fondue* methods. *get-sequences* method can be used to fetch raw sequencing data (single- and/or paired-end), while *get-metadata* can download corresponding metadata. Both methods can be run simultaneously by using the *get-all* pipeline. Sequences obtained from multiple fetches can be combined using *combine-seqs* and multiple metadata artifacts can be merged with *merge-metadata*. Run and Bioproject IDs can be retrieved from Zotero web library collections with the *scrape-collection* action.

All data-fetching actions support configurable parallelization to maximally reduce the processing time. The *get-metadata* method employs a multi-threading approach built into the *Entrezpy* modules (see the *Metadata retrieval* section), while *get-sequences* uses multiple processes and queues to coordinate the data download with its pre- and post-processing steps.

#### Sequence retrieval

The *get-sequences* action makes use of the *sra-tools* command-line toolkit (Leinonen, Sugawara, et al., 2011) https://github.com/ncbi/sra-tools) developed by NCBI. The *prefetch* tool is first used to reliably fetch SRA datafiles using the provided run IDs and the *fasterq-dump* utility is then executed to retrieve the corresponding sequences (single- or paired-end) in the FASTQ format. To follow QIIME 2’s naming convention those sequences are then renamed using their accession IDs, compressed, and finally validated by QIIME 2’s built-in type validation system. The action keeps track of any errors that occurred while fetching the sequences and performs available storage checks on every iteration to ensure no data is lost when space is exhausted during download. There are three output files generated by the *get-sequences* action: two QIIME artifacts corresponding to single- and paired-end reads, respectively, and one table containing the list of IDs for which the download failed (if any) including the linked error messages.

#### Metadata retrieval

Retrieval of SRA metadata is possible through the *get-metadata* action. This action uses the *Entrezpy* package (Buchmann & Holmes, 2019) to interact with the SRA database by building on top of its built-in modules for different E-Direct utilities. Specifically, we implemented a new *EFetchResult* and *EFetchAnalyzer* that work in tandem to request and parse metadata for the provided run IDs. The result is represented as a table where a single row corresponds to one SRA run and columns reflect all the metadata fields found in the obtained response. In order to keep track of different metadata levels (study, sample, experiment, and run), we introduced a set of cascading Python data classes to delineate the hierarchical relationship of the SRA metadata entries (Figure 3) and to preserve this structure in the final study metadata. Moreover, tight integration with QIIME 2’s internal type validation system guarantees consistency of metadata generated by *q2-fondue* by ensuring presence of all required metadata fields, as specified by NCBI.

**Figure 3.**
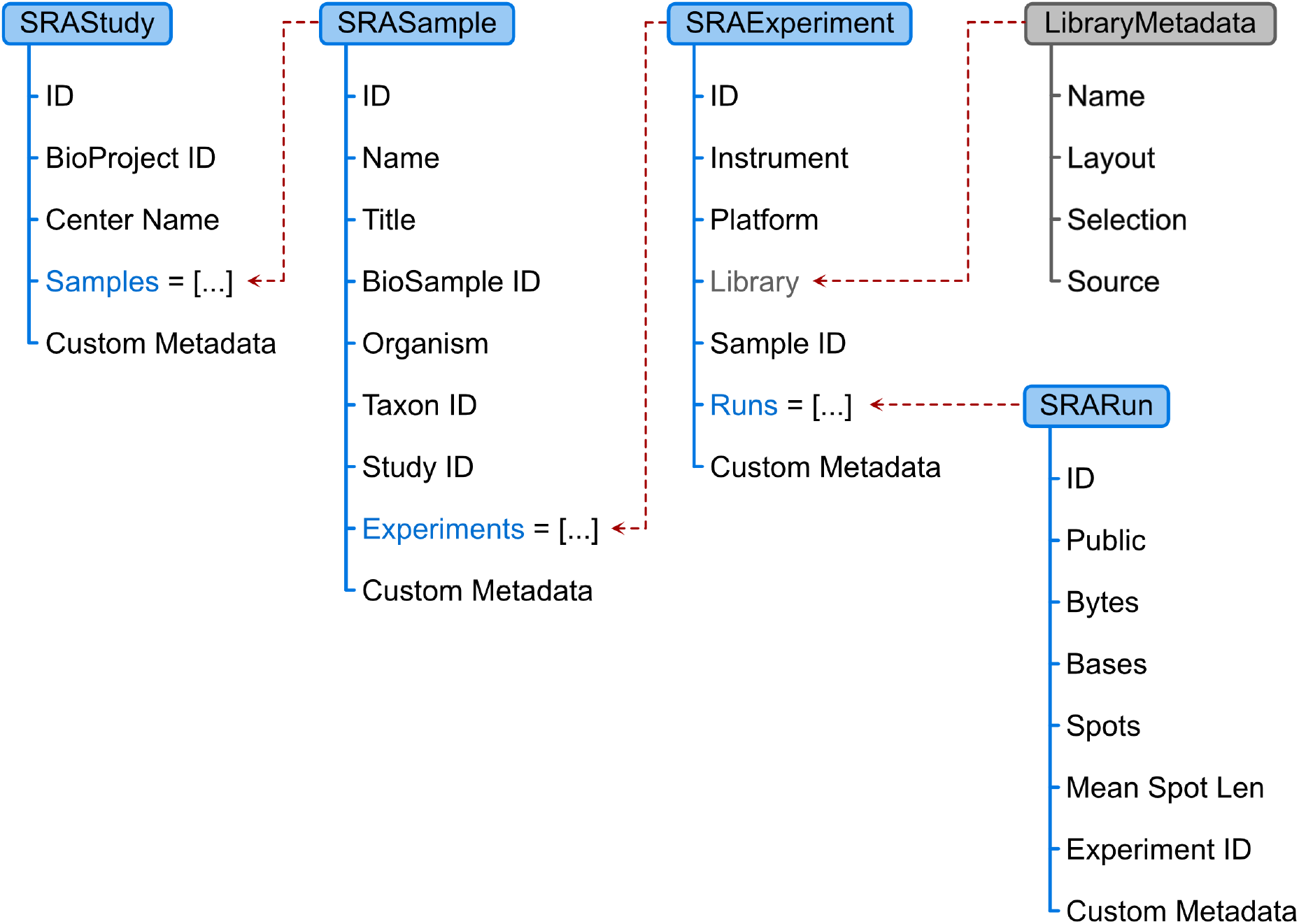
Structure of the SRA metadata *data classes* used by the *get-metadata* action. Each of the classes represents a different level of metadata organization and can contain other nested metadata objects. As all the objects are linked together, metadata of the entire study and all its children can be retrieved directly from the SRAStudy object.

#### Other available actions

The q2-fondue package contains additional functions to simplify sequencing (meta)data retrieval and manipulation, particularly when multiple data fetches are necessary.

➔ ***get-all*** allows simultaneous download of sequences and related metadata.
➔ ***merge-metadata*** concatenates metadata tables obtained from several independent *get-metadata* runs and allows generation of a single, unified metadata artifact.
➔ ***combine-seqs*** merges sequences obtained from multiple artifacts obtained from several *get-sequences* runs (or from other external sources) into a single sequence artifact.
➔ ***scrape-collection*** retrieves SRA run and BioProject IDs from a Zotero web library collection (https://www.zotero.org) by using the *pyzotero* package (Hügel et al., 2019). This enables seamless workflows for collecting IDs of interest from a literature collection, automatically downloading the data, and processing downstream with q2-fondue and QIIME 2.

### Use cases

The *q2-fondue* plugin can be used to facilitate comparative analysis of any nucleotide sequence data and metadata available on the SRA. To demonstrate some example use cases, we used *q2-fondue* and QIIME 2 to analyze marker-gene, whole genome sequence, and shotgun metagenome data. All analyses described below are available as fully reproducible and executable Jupyter notebooks (available in Supplementary material). These examples are provided merely as method demonstrations to showcase seamless integration/interoperation of q2-fondue with downstream analyses, and do not represent complete analyses of biological data. Additional steps and larger comparative analyses would be required to derive meaningful conclusions.

#### Marker-gene amplicon sequence data analysis

Marker-gene amplicon sequencing (e.g., of 16S rRNA genes) is currently the most popular method for high-throughput, untargeted profiling of microbial communities. To demonstrate the use of *q2-fondue* for comparative cross-study analysis of marker-gene amplicon sequence data, we selected three studies that analyzed the development of the infant gut microbiome in distinct geographical locations: (Lewis et al., 2017) with BioProjectID PRJEB16321, (Davis et al., 2017) with BioProjectID PRJEB15633 and (McClorry et al., 2018) with BioProjectID PRJEB23239. All three studies sequenced 16S rRNA genes in the V4 region with the forward 515F primer and a read length of 251 to 253 nucleotides. (McClorry et al., 2018) additionally used the reverse primer R806.

The *get-all* action of *q2-fondue* was used to retrieve metadata and sequence data of all three studies from the SRA. To normalize metadata features of interest across studies, namely age and health status, we employed the Python library *pandas* (McKinney, 2010; Reback et al., 2020). The downloaded single-read gene sequences were filtered according to availability of metadata with the *q2-demux* QIIME 2 plugin (https://github.com/qiime2/q2-demux), denoised with the *q2-dada2* QIIME 2 plugin (Callahan et al., 2016), https://github.com/qiime2/q2-dada2) and finally features were filtered by frequency, rarefied and summarized with the *q2-feature-table* QIIME 2 plugin (N. A. Bokulich, Kaehler, et al., 2018) https://github.com/qiime2/q2-feature-table). Finally, the processed metadata and sequence data were used to train two Random Forest classifiers with *q2-sample-classifier* (N. A. Bokulich, Dillon, et al., 2018), https://github.com/qiime2/q2-sample-classifier) to predict an infant’s age and its health status from its gut microbiome. The infant’s age was reported in four binned age groups: 0-1, 1-4, 4-6 and older than 6 months. For the health status, an infant was identified as healthy if no disease-related features were reported, namely no indication of stunting, wasting, underweight, elevated C-reactive protein status, parasites or anemia.

Both classifiers were trained and tested with 10-fold cross-validation using Random Forest classifiers grown with 500 trees. The performance of the trained classifiers was evaluated on the area under curve (AUC) of the Receiver Operating Characteristics (ROC) curve of the test set of each fold using *scikit-learn* implementations (Pedregosa et al., 2011). The entire analysis can be reproduced by executing the *u1-amplicon.ipynb* Jupyter notebook, available in the Supplementary material.

#### Comparative whole-genome sequence data analysis

The initial list of SARS-CoV-2 whole genome sequencing data accession IDs was generated based on the metadata obtained from the Nextstrain.org platform ((Hadfield et al., 2018); https://data.nextstrain.org/files/ncov/open/metadata.tsv.gz, access date: 08 February 2022). Only records with available SRA accession IDs and the least amount of missing data (column *QC_missing_data == ‘good’*) were retained. Furthermore, to limit the scope of this use case only Alpha, Delta and Omicron variants of SARS-CoV-2 virus were considered and all the Nextstrain clades corresponding to those variants were collapsed into three groups (*Nextstrain_clade_grouped* column). The *get-metadata* action of *q2-fondue* was used to retrieve metadata of 112’500 randomly sampled genomes (37’500 genomes per variant) from the SRA. Obtained metadata was then supplemented with the original Nextstrain metadata by merging the two datasets on common SRA run ID and only samples sequenced using single-end reads on the Illumina platform were retained. Finally, sequencing data for 750 randomly sampled genomes (250 genomes per variant) was fetched from the SRA using *q2-fondue’s get-sequences* method. Reads shorter than 35 nt were discarded using the *trim_single* method from the *q2-cutadapt* plugin ((Martin, 2011); https://github.com/qiime2/q2-cutadapt) and quality of the sequences was evaluated using the *summarize* action from the *q2-demux* QIIME 2 plugin (https://github.com/qiime2/q2-demux). MinHash signatures were computed for every sample and compared using the *q2-sourmash* plugin (*compute* and *compare* methods, respectively) ((Ondov et al., 2016); https://github.com/dib-lab/q2-sourmash). A t-SNE plot was generated from the resulting distance matrix using *q2-diversity (tsne* method with a learning rate of 125 and perplexity set to 18, https://github.com/qiime2/q2-diversity, (Halko et al., 2011)) and visualized using *matplotlib* and *seaborn* Python packages (Hunter, 2007; Waskom, 2021). Finally, to determine whether MinHash signatures are predictive of SARS-CoV-2 genomic variants, k-nearest-neighbors classification with 10-fold cross-validation was applied through the *q2-sample-classifier* plugin ((N. A. Bokulich, Dillon, et al., 2018); https://github.com/qiime2/q2-sample-classifier). The entire analysis can be reproduced by executing the *u2-genome.ipynb* Jupyter notebook, available in the Supplementary material.

#### Shotgun metagenome sequence data analysis

To fetch metadata for all the shotgun metagenome samples from the Tara Oceans Expedition (Tara Oceans Consortium Coordinators et al., 2015) we used *q2-fondue’s get-metadata* action with the following BioProject IDs: PRJEB1787, PRJEB4352, PRJEB4419, PRJEB9691, PRJEB9740, PRJEB9742. After removal of missing values, the resulting metadata table was randomly sampled to 100 records and used as input to the *draw-interactive-map* action from the *q2-coordinates* plugin ((N. Bokulich & Caporaso, 2018); https://github.com/bokulich-lab/q2-coordinates) to visualize values of the sensors used during the expedition according to their geographical location. Additionally, sequences corresponding to 10 samples at two different locations were fetched using the *get-sequences* action. To reduce the computational time required for this use case demonstration, the reads were subsampled to 20% of the original read count and the resulting artifact (containing single-end reads) was used to calculate and compare MinHash signatures (see previous section). A PCoA analysis was performed on the resulting distance matrix using the *pcoa* action from the *q2-diversity* plugin (https://github.com/qiime2/q2-diversity, (Halko et al., 2011)) and the PCoA plot was visualized using *matplotlib* and *seaborn* Python packages (Hunter, 2007; Waskom, 2021). The entire analysis can be reproduced by executing the *u3-metagenome.ipynb* Jupyter notebook, available in the Supplementary material.

## Results

Any meta-analysis can be carried out using raw experimental data, its associated metadata, or a combination of both. To demonstrate the versatility of *q2-fondue* in all those scenarios, and seamless integration/interoperability with downstream bioinformatics tools, we performed three example use case meta-analyses using amplicon, whole genome, and shotgun metagenome sequencing data and related metadata. All three use cases exclusively employ QIIME 2 plugins to process received data, and illustrate how *q2-fondue* can immensely increase data analysis reproducibility and transparency by including details on the raw data fetching in the QIIME 2 provenance.

### Use case 1: amplicon sequencing

As a demonstration of *q2-fondue*’s capacities in enabling the collection and analysis of amplicon sequencing data, we selected three infant gut microbiome development studies from distinct geographical locations: the study by (Lewis et al., 2017) from Georgia, (Davis et al., 2017)’s study from Gambia and (McClorry et al., 2018)’s study from Peru. We used the BioProject IDs reported by those studies to fetch the corresponding raw metadata and sequencing data. This provided us with 350 sequence samples each annotated with 148 metadata features.

After performing filtering, normalization, and denoising steps on the raw 16S rRNA gene sequences (see Figure 4A for an overview of plugins and actions used throughout this use case), a total of 3’880 amplicon sequencing variants (ASVs) were identified for 330 samples. The available metadata was used to define binned age groups. The distribution of samples per age group as well as the analyzed age range differ per study (Figure 4B). We further defined a binary health status which denotes whether the sample stems from a healthy or unhealthy infant (see Methods for more details). Across all studies, 194 unhealthy and 136 healthy infant samples were identified. Figure 4C displays the fraction of healthy infants in each of the three geographic locations covered by the selected studies.

**Figure 4.**
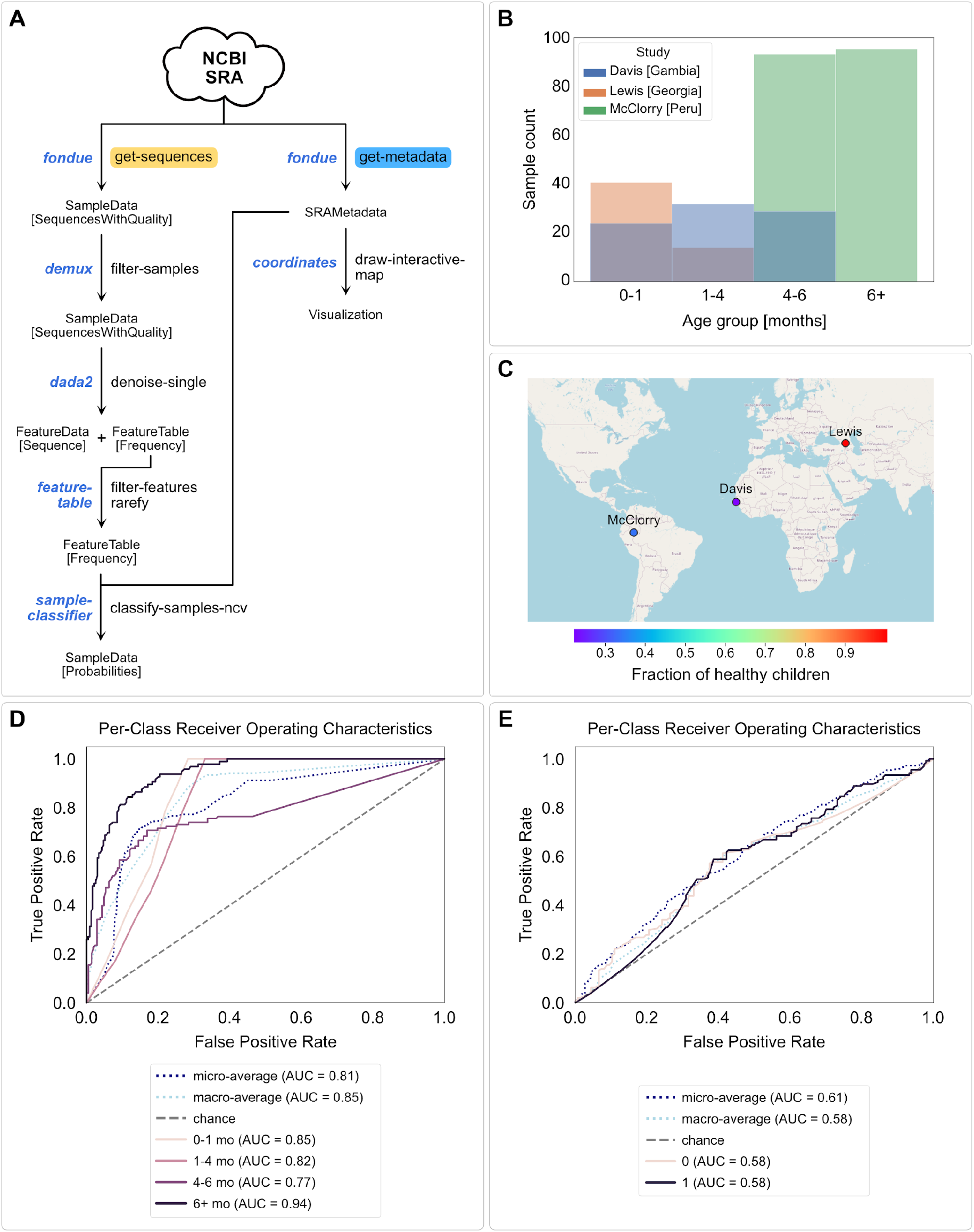
Analysis of amplicon sequencing data from three infant gut microbiome development studies. (**A**) Overview of QIIME 2 actions used during the amplicon data analysis. (**B**) Counts of samples in the defined age groups per study. (**C**) Fraction of healthy infants in the geographic locations covered by the selected studies. (**D** & **E**) ROC curves of Random Forest classifiers predicting age groups (D) and health status (E), indicating better predictive accuracy for age groups (macro averaged AUC = 0.85) than health status (macro averaged AUC = 0.58).

Finally, we trained two Random Forest classifiers with 10-fold cross-validation on the processed microbiome sequence data to predict age group and health status of each sample, respectively. The classifiers were evaluated on the test set of each fold and revealed a better performance in predicting age groups (macro averaged AUC = 0.85, Figure 4D) than health status (macro averaged AUC = 0.58, Figure 4E).

### Use case 2: whole genome sequencing

To illustrate how *q2-fondue* can be used as an entry-point to analysis of whole genome sequencing data we turned to one of the most rapidly growing datasets of the recent years: the SARS-CoV-2 genome dataset. We used all of the pre-processed metadata obtained through the Nextstrain.org platform (Hadfield et al., 2018) to identify samples which have been deposited in the SRA. We subsampled genomes of three SARS-CoV-2 variants: Alpha, Delta and Omicron (according to Nextstrain’s clade naming strategy and WHO’s variant labeling convention). We then fetched the corresponding SRA metadata using *q2-fondue*, which was used to prepare our final list of genomes. To simplify the analysis and reduce technical variability, we focused only on samples sequenced using single-end reads on the Illumina NextSeq 550 platform. Following the quality control step, we used the *sourmash* tool to readily compare viral genomes to one another by computing their MinHash signatures (Ondov et al., 2016). The resulting distance matrix was then used to generate a t-SNE plot visualizing how sampled genomes group together. Figure 5B shows that the Omicron variant forms a separate cluster in the t-SNE space and the Alpha and Delta clades, while distinguishable from one another in a form of several smaller clusters, cannot be as clearly separated into two groups. Finally, we used k-nearest-neighbors clustering to quantitatively compare genome MinHash signatures to predict SARS-CoV-2 clade membership (Figure 5C). We found that it was possible to classify the three SARS-CoV-2 variants with an accuracy of 93%. An overview of plugins and actions applied in this use case can be found in Figure 5A.

**Figure 5.**
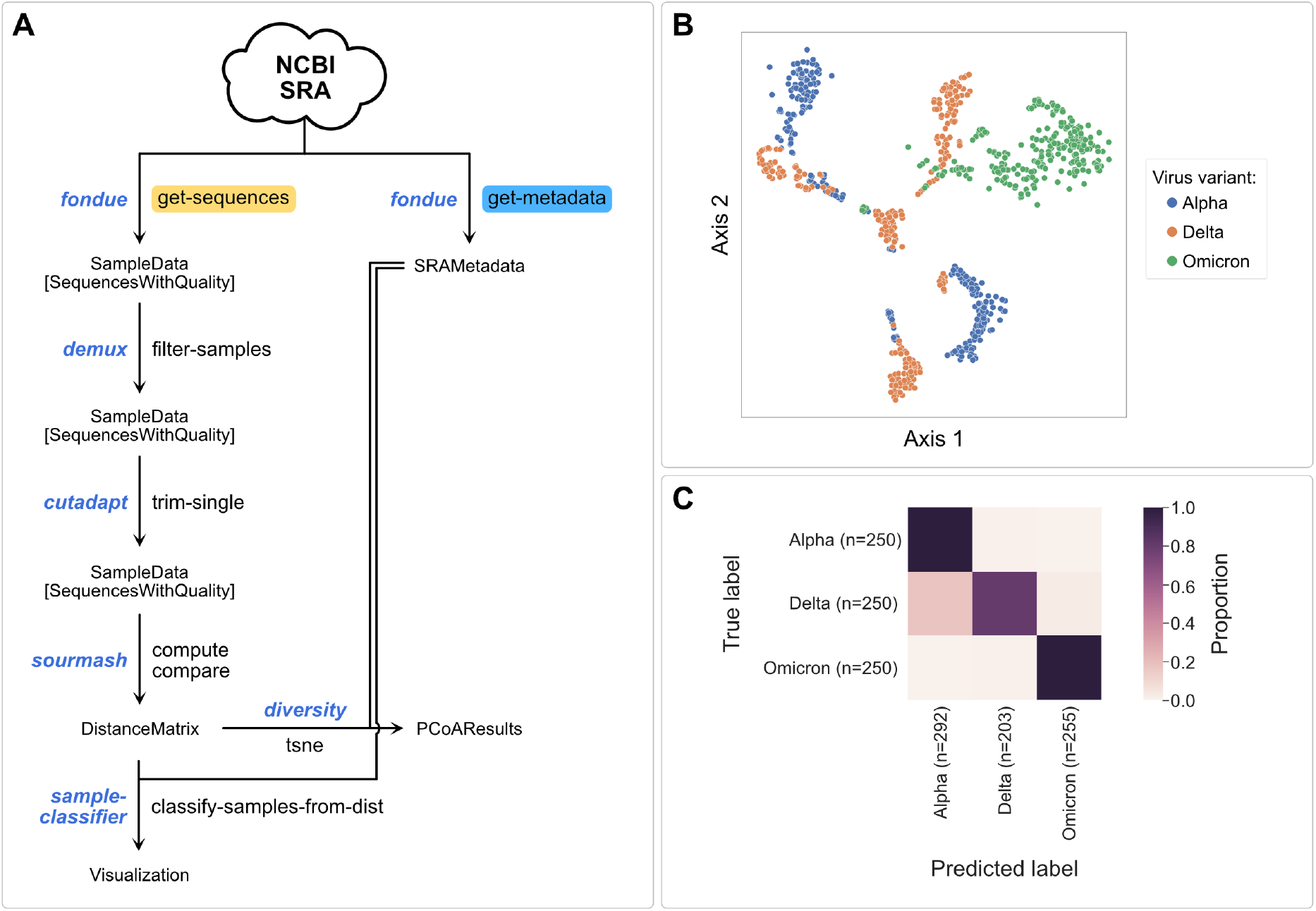
Genome MinHash signatures are predictive of SARS-CoV-2 variant type. (**A**) Overview of QIIME 2 actions used during the genome data analysis. (**B**) t-SNE analysis of the SARS-CoV-2 genome MinHash distance matrix shows that virus variants can be grouped into distinct clusters based on only genome hash signatures. (**C**) The same distance matrix can be used to reliably predict virus clades from genome hashes. K-nearest-neighbors clustering approach with 10-fold cross-validation was used to classify samples - a confusion matrix constructed from all test sets is shown.

### Use case 3: shotgun metagenome sequencing

We used the Tara Ocean expedition dataset (Tara Oceans Consortium Coordinators et al., 2015) to illustrate how geographic location included in sample metadata deposited in the SRA can be used to display sample properties, using *q2-fondue* and QIIME 2 (see Figure 6A for an overview of plugins and actions used throughout this use case). We fetched metadata for six BioProjects containing 1’049 ocean samples obtained through size fractionation followed by shotgun metagenome sequencing (Figure 6B-C). As geographical coordinates of every sample are included in the SRA metadata, we could directly draw an array of interactive maps visualizing various sample properties using the *q2-coordinates* plugin (N. Bokulich & Caporaso, 2018). As an example, Figure 6D illustrates sample temperatures across the globe. Moreover, we randomly selected 10 samples collected at two distinct locations and used the corresponding sequences to calculate and compare their MinHash signatures. Using PCoA analysis of the resulting distance matrix, we could show that the samples can be separated by location when using only their genome hash signatures (Figure 6E). More interactive visualizations can be found in the Jupyter notebook accompanying this manuscript (see Supplementary material).

**Figure 6.**
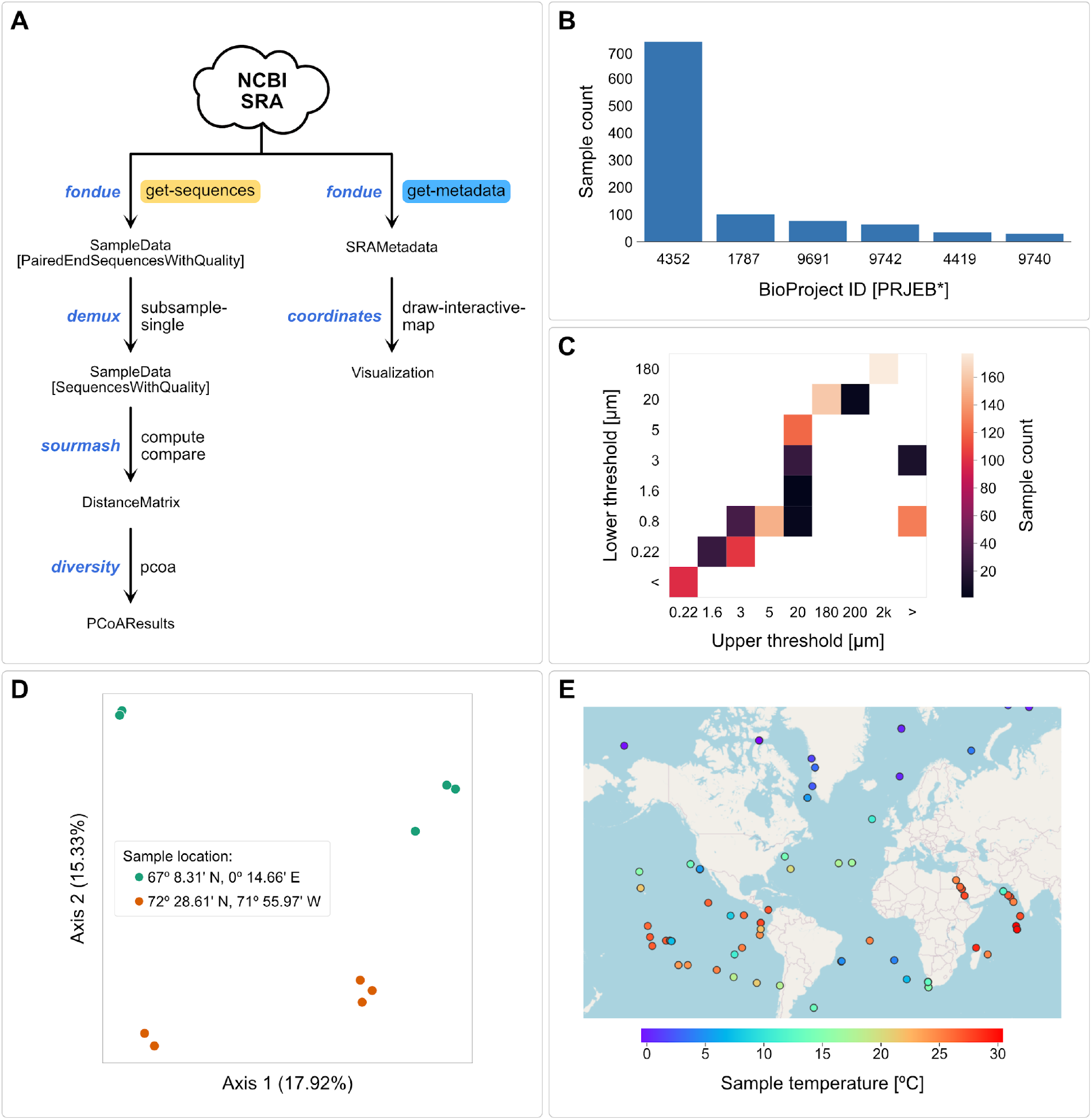
Tara Oceans expedition (meta)data analysis. (**A**) Overview of QIIME 2 actions used during the metagenome analysis. (**B**) Counts of samples in the retrieved dataset according to BioProject ID. (**C**) Counts of samples corresponding to different fractions obtained through size fractionation. (**D**) PCoA analysis of the metagenome MinHash signatures of 10 samples taken at two randomly selected locations. Fraction of explained variance is shown for two plotted dimensions. (**E**) Temperature of samples taken at different geographical locations. Only 100 randomly selected samples are shown.

### Integration with QIIME 2 ecosystem

Since *q2-fondue* is a QIIME 2 plugin, it tightly integrates with and benefits from the rest of the QIIME 2 ecosystem. Sequences obtained through the *get-sequences* action can be directly plugged into any other QIIME 2 action that operates on this data type (see Figure 7 for an overview of actions applied in this study). In addition to defining format checks for SRA metadata objects, *q2-fondue* has implemented transformer functions to allow the metadata downloaded through the use of *get-metadata* action to serve as input to any QIIME 2 action that requires sample metadata. Furthermore, integration with QIIME 2’s built-in provenance tracking system ensures that data fetching from the SRA is also included in the provenance graph (stored directly in all data outputs), enabling researchers to track and completely reproduce the entire analysis pipeline from data download to final visualizations.

**Figure 7.**
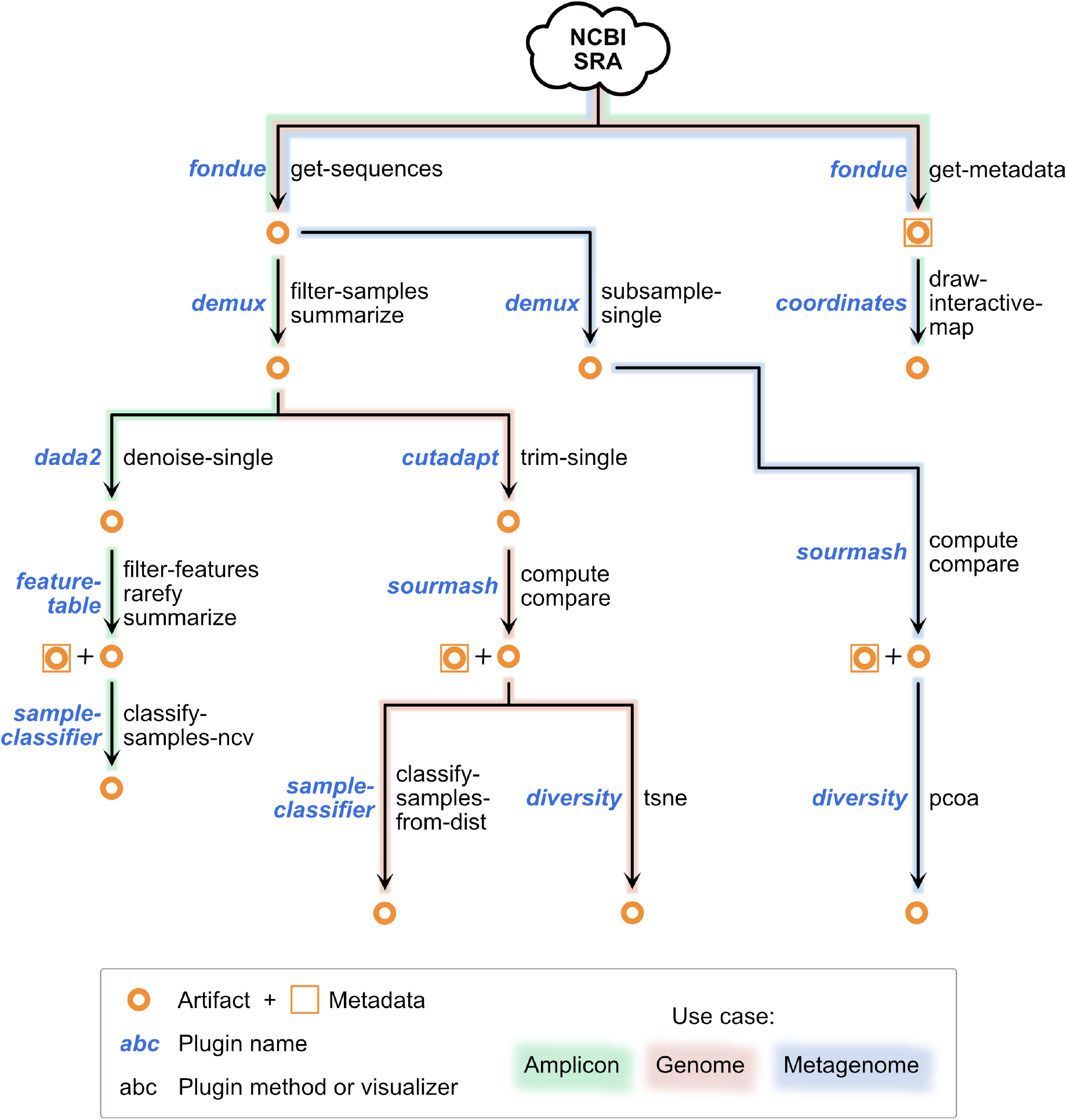
Overview of *q2-fondue* integration with other QIIME 2 plugins and actions as applied in the three use cases presented in this study. This is only a limited demonstration of possible downstream uses for three different nucleotide sequence data types, not an exhaustive list.

## Discussion

Declining costs and increasing throughput of nucleotide sequencing have fueled an exponential increase in published sequence data over the past two decades (Stephens et al., 2015). These data have an immense reuse potential, which has led to a growing trend of sequencing-based meta-analyses (Figure 1), paving the way to additional discoveries regarding general biological trends (Abbas et al., 2019; Panagiotou et al., 2013; Thompson et al., 2017). However, such studies remain technically challenging and data acquisition and management are significant bottlenecks. We developed *q2-fondue* to lower these hurdles, and to facilitate reproducible acquisition and management of metadata and nucleotide sequence datasets from the SRA (see Table 1 for a summary of the most important features). Its integration with the QIIME 2 framework offers complete provenance tracking of the entire process, multiple user interfaces, and thorough input/output data validation, allowing to conduct meta-analyses in a reproducible manner. Furthermore, *q2-fondue* outputs can be directly used with a wide range of QIIME 2 plugins, offering the user a smooth incorporation with any sequence-based analysis that is (or will become) available within the QIIME 2 ecosystem. Finally, *q2-fondue*’s integration with QIIME 2 offers users unparalleled support through the QIIME 2 forum - an exchange platform between users and plugin developers (with a current total of 5’700 signed-up members).

**Table 1.** Selection of the most significant issues faced by users when retrieving large amounts of sequencing data, together with their user-friendly solutions offered by *q2-fondue*.

| Problem | Solution offered by *q2-fondue* |
| --- | --- |
| Plethora of accession ID types complicates retrieval of sequences/metadata. | Conversion between BioProject and run IDs is performed automatically. All possible accession IDs are recorded in the final metadata table. |
| Potential data loss on space exhaustion when fetching large amounts of runs. | *q2-fondue* keeps track of available disk space and will abort without data loss when the amount of space is insufficient. |
| Sequencing data requires preprocessing/name normalization before it can be used in downstream analyses. | *q2-fondue* takes care of renaming/standardizing all the files after retrieval. |
| Merged datasets and subsequent data analysis steps are not always reproducible. | Tight integration with QIIME 2 ensures that every data fetching and analysis detail is recorded in provenance stored together with every single output. |
| Diversity of metadata fetched from multiple studies complicates its application in subsequent analyses. | Metadata retrieved by *q2-fondue* is normalized into a single table with standardized columns. |
| Network issues and other errors lead to data loss and require cumbersome, repeated data fetches. | Data retrieval can be automatically repeated in case of encountered errors. In case of repeated failures, all errors are reported and can be investigated by the user once the download is finished. No data loss occurs. |
| Parallelization of custom SRA access scripts is complicated and time-consuming. | *q2-fondue* takes care of data retrieval/processing in a parallel way, making use of multiple threads and CPUs available on the user's system. |

Despite its ease of use, *q2-fondue* does not free the user of their due diligence in checking the details on the extracted datasets in the accompanying publications, where mismatches with obtained run metadata or sequences could be detected.

The *q2-fondue* demonstrations shown here represent only a few possible use cases for the software, and we envision many other possible applications for analysis of diverse nucleotide sequence data types.

### Ideas and Speculation

The *q2-fondue* package remains under active development, and several additional functionality upgrades are planned in the future. As *q2-fondue* operates on large amounts of sequencing data we will introduce several performance-enhancing updates that will allow better management of free storage space available during download as well as streamline downloading large numbers of accession IDs to avoid multiple re-fetches.

While using run and BioProject accession IDs may cover needs of a large group of users, we are planning to additionally enable retrieving data using other kinds of IDs, notably SRA’s Study ID, such that even more studies deposited in SRA can be downloaded using *q2-fondue* in a more flexible way.

*q2-fondue*’s metadata retrieval action already greatly simplifies downloading metadata of multiple projects and formatting those as a single result table. Several additional functions are planned to assist with management and integration of diverse study and sample metadata.

Finally, to unlock the potential of sequencing data stored in and processed by other repositories we will add support for (meta)data retrieval from various other databases (e.g., MGnify (Mitchell et al., 2020)). Altogether, we hope that *q2-fondue* can become the tool of choice for interacting with SRA and other similar repositories, while at the same time seamlessly integrating with the whole QIIME 2 ecosystem, hence enabling a wide range of available analysis types.

The plugin is licensed under BSD-3-clause license and available under https://github.com/bokulich-lab/q2-fondue.

## Supplementary material

An introductory tutorial for the usage of *q2-fondue* is available under https://github.com/bokulich-lab/q2-fondue/blob/main/tutorial/tutorial.md. All three use cases can be reproduced by executing the respective Jupyter notebooks accompanying this article (https://github.com/bokulich-lab/q2-fondue-examples).

## Acknowledgements

We thank Evan Bolyen (Northern Arizona University) for insightful discussions on metadata processing and working with the SRA and SRA Toolkit. We also thank Anton Lavrinienko (ETH Zürich) for his valuable comments on the manuscript.

